# Tuning Ultrasensitivity in Genetic Logic Gates using Antisense RNA Feedback

**DOI:** 10.1101/2024.07.03.601968

**Authors:** Nicolai Engelmann, Maik Molderings, Heinz Koeppl

## Abstract

This work provides a study of a possible improvement of existing inverting genetic logic gates by introduction of a common sequestration reaction between their input and output chemical species. As a mechanism of study, we use antisense RNAs (asRNAs). The asRNAs are expressed with the existing messenger RNA (mRNA) of a logic gate in a single transcript and target mRNAs of adjacent gates, creating a feedback of the protein-mediated repression that implements the core function of the logic gates. The extended transcripts then share a common sequestration reaction mediated by the cellular host’s RNA metabolism. This sequestration consists of double-stranded RNA (dsRNA) formation by asRNA and adjacent mRNA and subsequent degradation by the host. Numerical and stochastic analysis suggests that the feedback increases the steepness of the gate’s transition region, reduces the leakage, and can potentially be used to adjust the transition location. To leverage these effects, we demonstrate how design parameters can be tuned to obtain desired dose-response curves and how arbitrary circuits can be assembled using the improved gates.

## 1 Introduction

The constituents of many cellular biochemical processes, such as the binding of regulator proteins to DNA binding-sites, appear to be related by dose-response curves that have sigmoid shape in a logarithmic argument (*1*). This phenomenon has enabled synthetic biologists to engineer artificial genetic pathways that compute logic operations *in vivo* (*2*). Competition for regulators by multiple binding domains, e.g. causing titration effects (*3*), can increase or decrease the steepness of dose-response curves in the logarithmic domain (*4, 5*). Strong binding affinity to competing sites can effectively lead to a sequestration of regulators (*6*), and it has been observed that sequestration influences dose-response curves in a similar way (*7*). Qualitative effects like this are of peak interest to researchers trying to engineer dose-response curves that behave like ideal logic switches (*8*). At the same time, recent interest in computer-aided design (CAD) of complex genetic logic functions has raised demand for highly modular and reusable devices (*9*). In Cello (*10, 11*), the most prominent of such CAD tools, logic computation is done by NOT and NOR gates only, implemented by proteins, specifically transcriptional repressors, and their corresponding promoters. Such protein-based circuits have been widely explored due to their ability to perform complex functions (*12*).

However, a significant limitation of those circuits are their large response time (*13, 14*) and substantial metabolic burden imposed on host cells. The expression of heterologous proteins demands considerable cellular resources, leading to a metabolic load that can disrupt normal cellular functions and reduce overall efficiency and viability of the host organism. Therefore, protein-based circuits limit the scalability and practical applicability in many contexts (*15* –*17*). In contrast, RNA-based circuits offer a promising alternative, characterized by faster response time and lower metabolic load. RNA molecules are produced and degraded more rapidly than proteins, allowing RNA-based circuits to propagate signals faster than their protein-based counterparts (*14, 18*). Moreover, the metabolic cost of expressing RNA molecules is significantly lower than expressing proteins, which involves production of amino acid chains and post-translational folding on top of its messenger RNA (mRNA) production. Several synthetic RNA switches have been developed and implemented in various organisms, showcasing the versatility and efficiency of RNA-based regulators (*19, 20*). Examples include small transcriptional activating RNAs (STAR) (*21*), CRISPRi (*22*), toehold (*23*) and its eukaryotic version eToehold (*24*) switches. These RNA-based elements can control gene expression precisely and dynamically, often with higher specifity and lower energy requirements then their protein counterparts. RNA molecules also play natural regulatory roles within cells, as seen with long non-coding RNAs (lncRNAs) (*25*) and microRNAs (miRNAs) (*26*), which participate in intricate regulatory networks. This natural predisposition for regulation makes RNA an attractive substrate for engineering synthetic feedback mechanisms. For instance, RNA-based integral feedback controllers using antisense RNA (asRNA) have been designed to maintain homeostasis within synthetic circuits, ensuring robust performance despite any perturbations of the system (*27*). This asRNA has high target specificity and has wide usage in gene silencing in therapeutic applications (*28, 29*). In these applications, non-coding RNA (ncRNA) that is complementary to regions of the target mRNA acting as asRNA is transfected into cells. The asRNA binds to the mRNA and forms double-strand RNA (dsRNA) that is rapidly degraded by the host as a security measure, e.g. against viral attacks (*30*). Prokaryotic and eukaryotic cells alike maintain mechanisms that allow rapid degradation of dsRNA, although they may work very differently.

The goal of this work is to study the use of the asRNA architecture to mutually degrade RNA species adjacent to the TF-mediated transcriptional repression that constitutes the central part of an inverting logic gate. In this way, we effectively obtain a sequestration of the regulating species of the inverting logic gate. Because the logic gate is inverting and the sequestration is mutual w.r.t. the regulating species of adjacent logic gates in a circuit, we expect an escalation of their input sensitivity and thus a steeper dose-response curve. Although we introduce a sequestration, the idea is somewhat comparable to that of a bistable toggle-switch (*2*), and similar hurdles w.r.t. parameter tuning are to be expected (*31*). In contrast to the toggle-switch, however, our approach can flexibly be configured due to RNA base-pairing implementing the sequestration (*32*). We also expect a leakage-reduction due to the sequestration of leaking mRNA. Our method can be used on logic gates that have stricly inverting parts on the level of RNA (*11, 33* –*35*)(the NOT gates in (*10*)), but not on those that lack such parts (e.g. the NOR gates in (*10*)).

In the following sections, we briefly describe the biochemical mechanism behind the RNA feedback architecture. Then, we investigate its impact on a simple NOT gate similar to those in (*10, 11*). We take a look at its dose-response curve, describe important qualitative properties and perform a stochastic and numerical study. We then provide a toolset to calibrate important kinetic parameters of the gate, and finally give an outlook on how such gates can be assembled together systematically to build arbitrary combinational logic circuits.

## 2 Results

### 2.1 RNA Feedback for Genetic Logic Gates

Over the past decades, many logic operations have been implemented in cells, as reviewed by (*36*). NOT and NOR gates are logic-complete, meaning they can be used to compute any logic operation (*37*). These gates are simple to design using transcriptional repressors, which has led to their widespread adaptation (*10, 11*). However, using only transcriptional repressors results in an on/off switch with a broad transition region, leading to variability where cells can be in a logically inconsistent state. Cells and organisms require switches that are distinctly on or off, with no intermediate states. To address this, nature has developed ultrasensitive switches (*38* –*40*). Steep ultrasensitive switches can be achieved using different mechanisms: blocking, sequestration, and displacement (*41*). Transcriptional repressor-based switches are limited in steepness due to the cooperativity which can be achieved. RNA, feedbacking the translational process, adds minimal metabolic burden to the circuit (*14, 21, 42*). We propose an RNA-mediated feedback mechanism using sequestration, which can be easily integrated with existing NOT/NOR logic. This RNA feedback mechanism involves using trans-acting antisense RNAs (asRNAs) adjacent to the coding mRNA of the output transcription factor (TF) in a single transcript. The start and stop codons surrounding the mRNA coding region ensure TF translation. The antisense region complements the coding region of the input TF’s transcript, as illustrated in Fig. 1. Watson-Crick pairing forms double-stranded RNA (dsRNA), repressing translation due to RNA-RNA duplex degradation by various mechanisms in different hosts (*43* –*45*) or by blocking ribosome binding (*46*).

**Figure 1:**
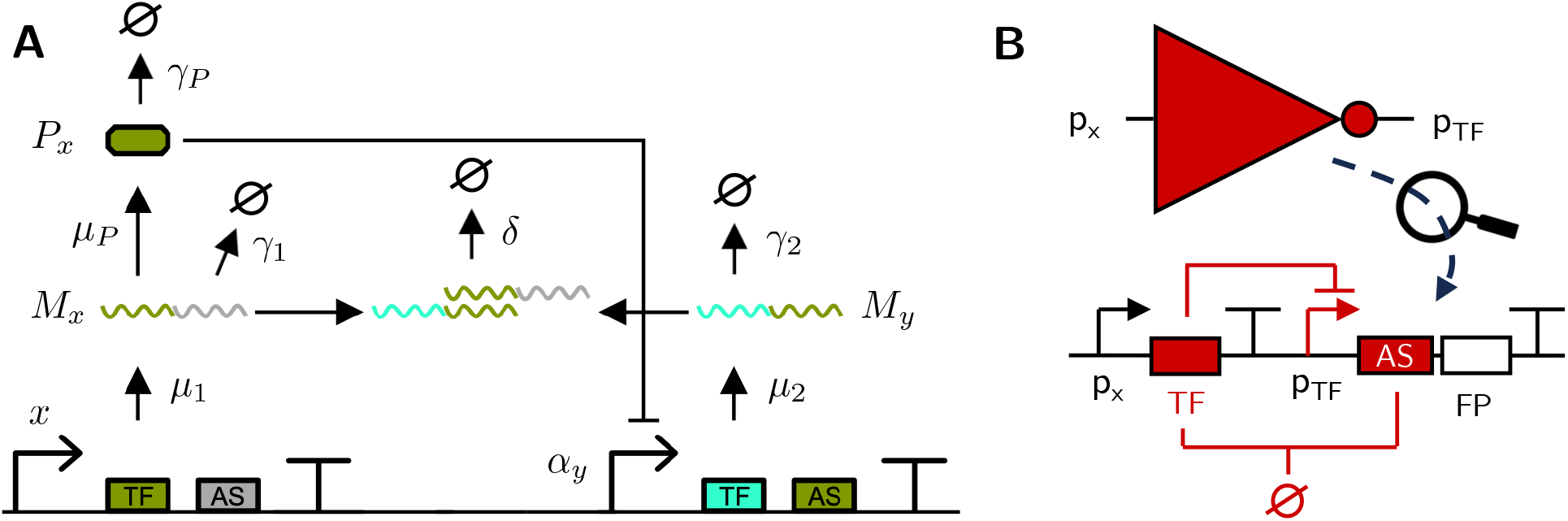
*A*) illustration of the proposed feedback architecture. Feedback is implemented through the transcription of RNA that includes both a coding and a non-coding segment. The coding segment is translated into a transcription factor, while the non-coding segment functions as an antisense RNA (asRNA) that targets the coding region of the input transcription factor. *B*) shows the SBOL diagram of an envisioned NOT gate with RNA-mediated feedback. The input species *x* is shown as a promoter activity and FP denotes a fluorescing protein as a potential output.

Inspired by regulation with antithetic integral feedback (*27, 47*), we propose that feedback mediation through sense and antisense RNA sequestration can enhance gate and circuit function. In eukaryotes, mRNA decay is relatively slow, whereas in prokaryotes like *E. coli*, mRNA decays within minutes (*48, 49*). To improve gate function in *E. coli*, dsRNA formation should occur within seconds. We hypothesize that the feedback is sufficiently fast because dsRNA formation by sense and antisense RNA in *E. coli* is a naturally occurring process (*50*), suggesting that dsRNA formation is faster than single-stranded mRNA decay.

### 2.2 The NOT Gate

In this section, the NOT gate – the most simple inverting logic element that can be extended by the RNA feedback connection – is considered. The SBOL diagram of the NOT gate is shown in Fig. 1B. It shares its structure with the NOT gate used in Cello (*10, 11*) with an additional asRNA as part of the transcript of the output chemical species. In the following, we analyse important features of the deterministic steady-state derived from the gate’s reaction rate equations and the given parameters of the original NOT gate without feedback.

#### 2.2.1 Steady-State Equations

Consider the reaction network of the simple NOT gate with feedback illustrated in Fig. 1A. It shows an arbitrary input species *X* that modifies the transcription rate of an RNA *M*_*x*_, which is then translated to protein *P*_*x*_. This protein acts as a repressing TF on the promoter that enables transcription of RNA *M*_*y*_. The quantity *α*_*y*_ denotes the promoter activity. *M*_*x*_ and *M*_*y*_ contain the matching sense and antisense regions that lead to dsRNA formation and successive degradation by the host’s responsible mechanism (e.g. RNase III in *E. coli* (*51*)). Small letters denote concentrations, e.g. *x ≡* [*X*] in the following. Note, that while *x* will most often represent the promoter activity of an input promoter to the gate taking values in a finite range, we can always multiply the RNA rate constant *µ*_1_ by a an arbitrary constant to rescale *x*. It thus makes sense for the scope of our analysis to work with a general input species *x ≥* 0. If not otherwise noted, we also generally assume a normalized reaction volume. The network’s reaction rate equations (RREs) are derived in Methods 4.1.2 using a quasi-steady state (QSS) assumption on the association and dissociation events of the TF *P*_*x*_ at the binding sites of the promoter. This simply allows us to treat *α*_*y*_ as a black-box Hill-function provided by the corresponding equation of the existing NOT gate. Note, that this is a common simplifying assumption (*52*). The steady-state equations can then be given by

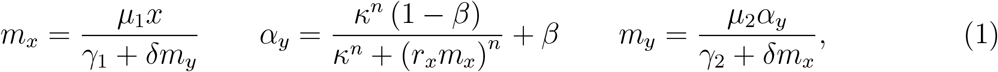

where *β ∈* [0, 1) determines the amount of leakage, 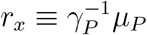 is a coefficient that generally depends on the inverter circuit used, and *n* is a Hill coefficient. Finding an explicit closed-form expression for *α*_*y*_ or *m*_*y*_ in *x* is not possible in the general case because of the feedback connection. However, numerical evaluation is possible, e.g. using a fixed-point iteration on (1) or solving a root-finding problem. These methods also scale to circuits of many gates with potential feedback connections (*6*).

#### 2.2.2 Limiting Behavior and Leakage Reduction

Maximal gene expression is reached for *x* = 0. Then, (1) suggests that *m*_*x*_ = 0, *p*_*x*_ = 0 and so *α*_*y*_ = 1 and in turn 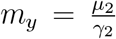. For the minimal expression limit, we let *x → ∞* and assume *m*_*y*_ *< ∞* stays finite. Then, *m*_*x*_ *→ ∞* becomes infinite and as a consequence *α*_*y*_ *→ β*. Hence, the concentration of the output RNA 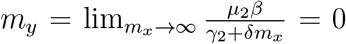 goes to zero. Clearly, *x* will be bounded in a realistic setting. However, in contrast to the original NOT gate with *δ* = 0, the steady-state leakage does not approach a constant asymptote for for *δ >* 0 with increasing *x*. This can be seen in Fig. 2C. Interestingly, a larger *δ* does not change the slope of the *asymptotic* decrease in *m*_*y*_ in a doubly logarithmic plot. This effect is not visible in Fig. 2C, but it comes from the circumstance that the promoter activity *α*_*y*_ saturates, while *m*_*y*_ has a hyperbolic relationship with *m*_*x*_ via the feedback. Assume *δ* and *x* large. Then, 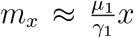 and *α*_*y*_ *≈ β*, and in turn 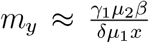. This is equivalent to 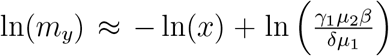. However, the point at which this asymptotic regime is reached varies with the value of *δ*, depending on when *γ*_1_*γ*_2_ + *δµ*_1_*x ≈ δµ*_1_*x* becomes valid. Note, that it is likely that in a real-world setting, a saturation of the leakage reduction by the feedback can occur not only due to the limited range of *x*, but also due to transport limitations (*53*) or additional cellular mechanisms. However, it is reasonable to assume that a reduction can be achieved to some degree.

**Figure 2:**
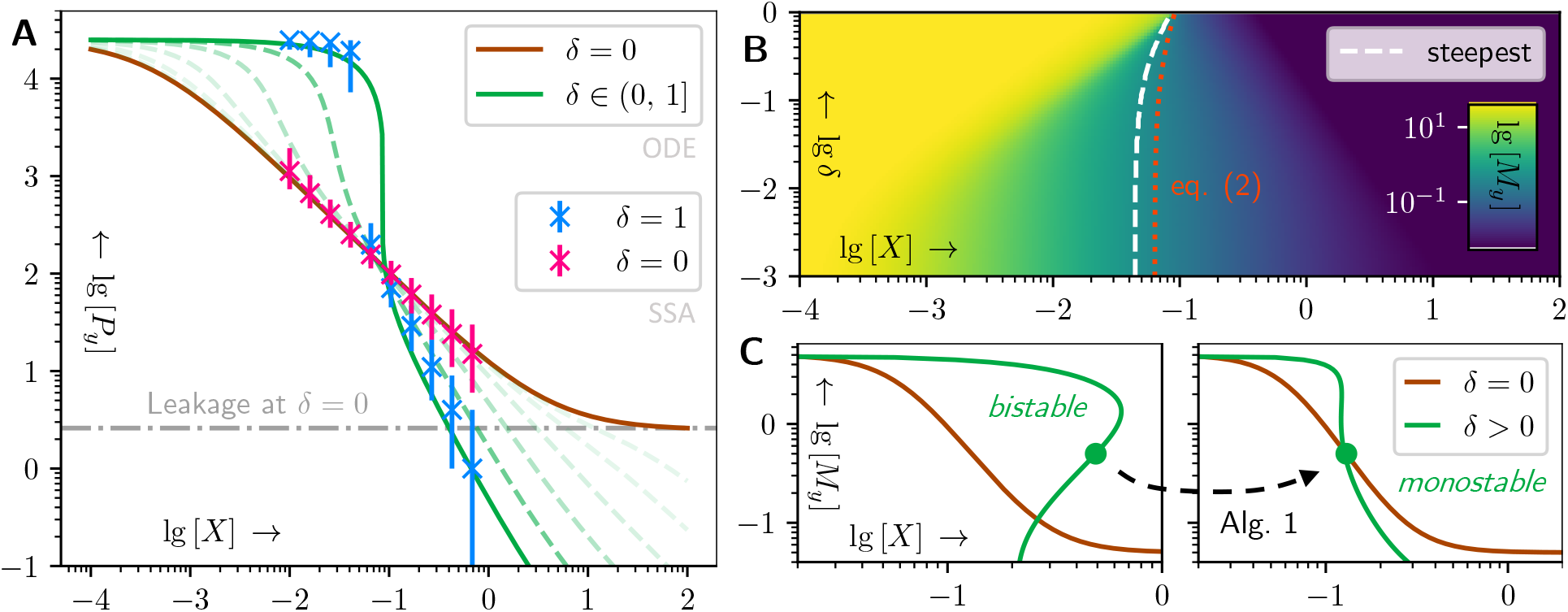
*A*) comparison of steady-state equations and stochastic simulation (SSA) of the exemplary NOT gate for different values of *δ*. The SSA *does not* employ the QSS assumption from the RREs. Details are found in Methods 4.1.1. *B*) shows the narrowing transition region with increasing *δ ∈* (0, 1] for the example NOT gate (c.f. Methods 4.1). The location of the steepest slope shifts in the process and the shape is no longer logistic. The reference location of the transition region *x*_tr_ from (2) is shown as a dotted red line. *C*) examplary usage of Alg. 1 for parameter tuning. The brown curves show the original dose-response. The green curves show steady-state curves (eq. (11) from Methods 4.1.3) and the marker shows the transition location *x*_tr_ (eq. (2)). Random initialization of *δ* can lead to bistability, Alg. 1 finds the optimal translation and sequestration rates *µ*_*P*_ and *δ*.

#### 2.2.3 Location and Uniqueness of the Logic Transition Region

The *logic transition region*, or simply the transition region, is a term from electrical engineering that denotes the region of a dose-response curve, in which the change between qualitatively distinct states mainly occurs (*54*). In a Hill-shaped dose-response curve, the transition region corresponds to the region around the inflection point, where the response changes rapidly with a change in dose. In this work, when we refer to the *location* of the transition region, we mean a specific reference dose that is deemed representative to the region. In a Hill-shaped dose-response curve, this could e.g. correspond to the dose at the inflection point.

In our NOT gate, the transition region in *m*_*y*_ becomes steeper the larger *δ* becomes, but also changes its location gradually with an increase in *δ* (c.f. Fig. 2B). The input species concentration *x* that causes the largest change in *m*_*y*_ can e.g. be found numerically from (1), iterating over *x*. However, locating the transition region by this criterion makes most sense when the dose-response curve is given by a Hill function because of its logistic shape in the logarithmic domain. This shape, however, is lost with *δ >* 0 and the dose-response curve exhibits no obvious symmetry. At the same time, the inflection point in the original Hill-shaped dose-response curve also corresponds to the input *x*, where the response has been attenuated by half the logarithmic difference between maximum and minimum response levels ln(*m*_*y*,max_) and ln(*m*_*y*,min_).

To obtain a useful criterion for all choices of *δ ≥* 0, we set the reference response level *m*_tr_ for the transition region to this value, i.e.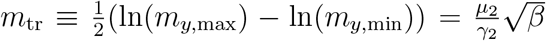, where *m*_*y*,max_ and *m*_*y*,min_ are the corresponding values of the original NOT case *δ* = 0. We can relate this choice to the concept of a threshold voltage in electronic logic gates (*54*). In Methods 4.1.3, we show that the input *x*_tr_ corresponding to *m*_tr_ can be obtained as the root-finding problem

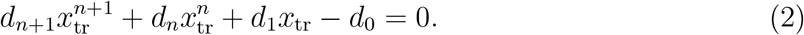

with the coefficients 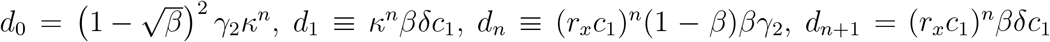, where we introduced the additional constant 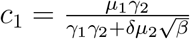. Because *β ∈* [0, 1], all coefficients are non-negative. Note, that the power *n* is not necessarily an integer. Equation (2) has been obtained by insertion of *m*_tr_ into the equation system (1) and subsequent elimination of *m*_*x*_ and *α*_*y*_. This does not lead to a loss of possible solutions for *x*_tr_, because given *m*_*y*_ = *m*_tr_, all equations in (1) are bijective in the remaining variables. Not only does (2) preserve all solutions, it has a unique non-negative real solution *x*_tr_ which we show in Methods 4.1.3. We plot *x*_tr_ as a function of *δ* for the example NOT gate in Fig. 2B as a red dotted line.

As a more general case, we also show in Methods 4.1.3 that there even exists a unique non-negative real input *x*_lv_ for any arbitrary *m*_lv_ *∈* [*m*_*y*,min_, *m*_*y*,max_]. This has the consequence that the extended NOT gate has a unique transition region in the important range [*m*_*y*,min_, *m*_*y*,max_]. We expect the dose-response curve w.r.t. *m*_*y*_ (and in consequence *a*_*y*_) to be monotonically decreasing with increasing *x* starting from an initial stable pair 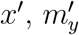, but a detailed analysis of this property is left open for future study.

Note, that we can have multiple *m*_tr_ that result in the same solution *x*_tr_ in (2), so that multiple steady-states of (1) can exist. Given the regulatory motifs in our construct, we can expect (1) to exhibit bistability for certain values of *δ*. While bistability of the dose-response curve akin to that of a bistable toggle-switch (*2*) or a Schmitt trigger (*55*) is not necessarily an unwanted effect in a switch, we show how to calibrate the sequestration rate *δ* to avoid this effect entirely in section 2.3.2.

### 2.3 Tuning Design Parameters

An existing NOT gate that is extended by the RNA feedback mechanism can exhibit a shift in its transition location and potentially loose its monostability. The latter effect can be desirable, as the presence of a hysteresis imposes a stabilizing effect on the set logic levels in the presence of fluctuations of the gate’s input. Still, a typical application would require most characteristic features of the original dose-response curve to stay unaltered, so that only its steepness increases and leakage decreases. At the same time, adjustments of design parameters to reach this goal must focus on those parameters that can be calibrated rather flexibly in the actual genetic design.

#### 2.3.1 Calibration of the Transition Location

Modification of the transition location *x*_tr_ can be done by changing the scaling coefficient 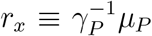 that controls the protein concentration [*P*_*x*_]. Tuning the translation efficiency of *P*_*x*_ e.g. by choosing an appropriate Kozak sequence or RBS, depending on the organism, implements the change in *r*_*x*_. Unfortunately, although *r*_*x*_ only appears in the equation for *α*_*y*_ in (1), it cannot simply be scaled by a factor 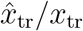 with the desired transition location 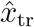 due to the feedback. However, the adjusted scaling coefficient 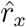 corresponding to the transition location 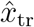 can be calculated explicitly by

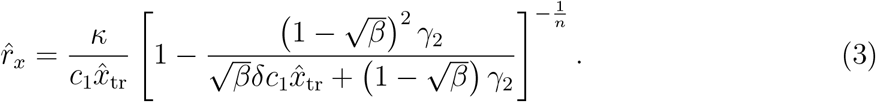

with the constant 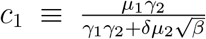. Eq. (3) can be obtained by rearranging (2) under knowledge that 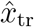 is unique.

#### 2.3.2 Preservation of Monostability

Monostability can be preserved by adjusting the sequestration rate *δ*. At the same time, *δ* can be tuned flexibly by modification of the hybridization strength between the engineered asRNA and its targeted mRNA. However, *δ* also determines the increase in steepness of the dose-response curve. Thus, a desirable *δ* would be as large as possible without causing multistability. To achieve this, we again consider (2) from section 2.2.3. We have shown in 4.1.3 that the LHS of (2) describes a function *f* (*x, m*_lv_) *∈ ℝ* that is strictly monotonically increasing in *x ∈* (0, *∞*) for constant *m*_lv_. Assume now, we have determined a root *x*_lv_ that satisfies *f* (*x*_lv_, *m*_lv_) = 0 for a specific *m*_lv_. Remember that a monostable inhibitory doseresponse curve is stricly monotonically decreasing in *m*_lv_ for increasing *x*_lv_. Thus, its inverse function is strictly monotonically decreasing in *x*_lv_ for increasing *m*_lv_ in its appropriate range.

We must therefore demand for any 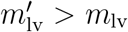 that the corresponding 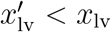 is smaller if both 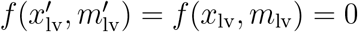. Now combining this with the monotonicity of 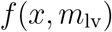 in *x* for any given 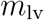, we must demand that 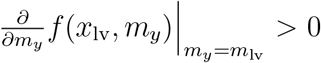 for any pair 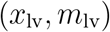 that satisfies 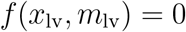.

This condition is hard to guarantee for all pairs 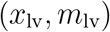. However, we can exploit the fact that the reference point for the transition location (*x*_tr_, *m*_tr_) conveniently lies in proximity to the presumed center of the region, where multistability may occur. We can thus propose the following heuristic that should yield satisfying results in most realistic cases: a good choice for *δ* is one, in which 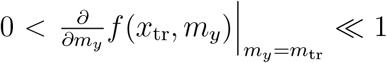 is satisfied. Note, that this partial derivative can be evaluated easily using automatic differentiators.

#### 2.3.3 The Combination of Both – Algorithmic Calibration of Parameters

Adjustment of the transition location and maintaining monostability is not as straightforward as simply applying the corrections from sections 2.3.1 and 2.3.2 one after another. Every change in *δ* causes the transition location *x*_tr_ to change and every adjustment of *r*_*x*_ reweighs the contributions of transcriptional repression and sequestration to the shape of the dose-response curve. However, it is possible to use an iterative approach to jointly calibrate *δ* and *r*_*x*_. An example algorithm that implements such an approach is given in Alg. 1 in Methods 4.2. We used Alg. 1 to find the optimal *r*_*x*_ and *δ* for the case shown in Fig. 2C, where *δ* has been initialized to a value that caused bistability and a significant shift in transition location. Using Alg. 1, we shifted the transition location back to its original value and chose a *δ* that was close to its largest possible value in the monostable regime. The initial and terminal parameters are given in Methods 4.2.

### 2.4 The Logic Circuit with RNA Feedback

In this section, we consider a full logic circuit whose gates are extended by RNA feedback. First, we investigate how an arbitrary existing gate architecture can be extended by RNA feedback and under which conditions. Second, we formally describe the full logic circuit with extended gates and present its steady-state equations.

#### 2.4.1 RNA feedback can extend Gates with Molecular NOR Logic

Many genetic gate architectures are composed of multiple parts that together implement the desired overall logic function. Given an existing architecture, a natural question to ask is how to properly integrate the RNA feedback mechanism in the architecture to obtain the desired effects and without disrupting the logic function. As an example, we can take a look at the NOR logic gates from (*11*). This specific gate architecture together with an extension by the RNA feedback is shown in Fig. 3B. Note, that the architecture in (*11*) uses gene copies with individual promoters instead of the tandem promoter architecture in (*10*). The latter is not compatible with the RNA feedback mechanism. This will be made clear in the following.

**Figure 3:**
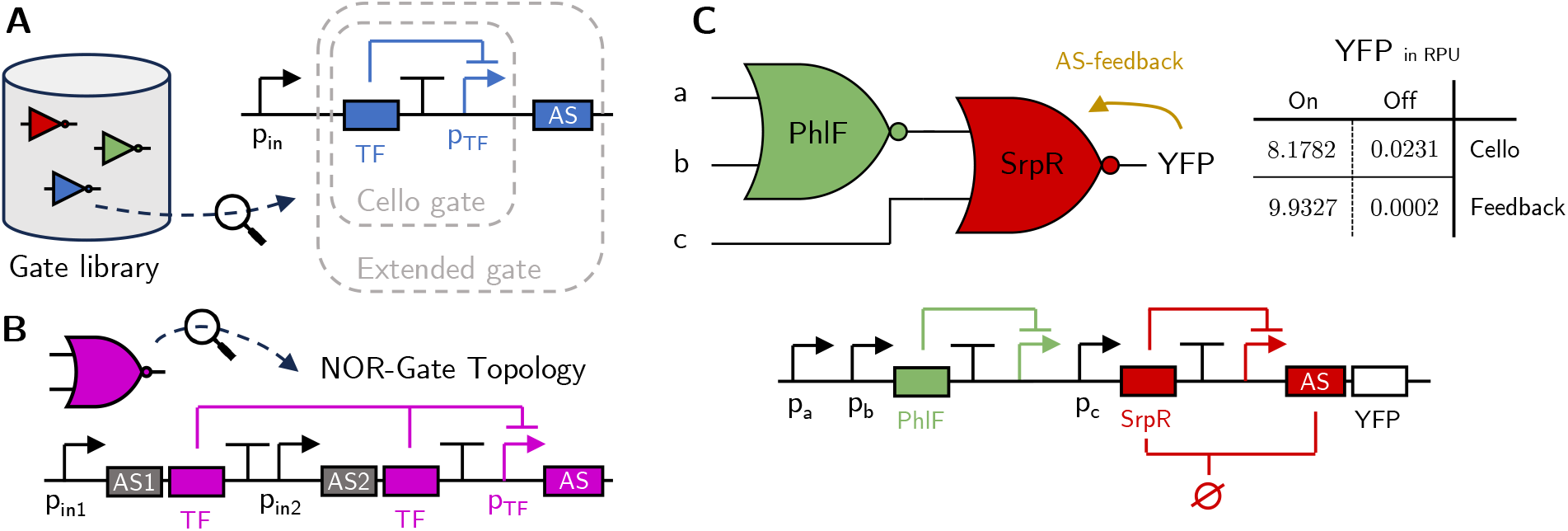
Construction of the NOR-logic circuit with RNA feedback. *A*) The genetic parts can be organized in a gate library. Compared to Cello’s gates, the extended gates contain an additional corresponding asRNA sequence that is placed next to the coding sequence of the output species. *B*) NOR gates can be assembled in accordance to the logic function using the strategy in (*11*), so that no mRNA is accidentally targeted that should exhibit a high copy-number. *C*) Cello circuit 0×70 with optimal gate assignment and RNA-feedback from YFP to SrpR; the table shows the lowest ON and highest OFF concentrations of YFP. We stress that this is an *in silico* experiment under the model assumptions made in this work.

Mutual repression of translation between species of logic gates can impact their logic function. Therefore, a gate with multiple inputs logically consistent with the single-transcript RNA feedback connection must implement a generalized inverter structure on the *molecular* level. Let *O ∈* {0, 1} be the Boolean output of an *N* -input logic gate and *I*_*n*_ *∈* {0, 1} for *n ∈ ℐ ⊂* ℕ its enumerated Boolean inputs. Let *T ⊂ ℐ* be the indices of the inputs targeted by RNA feedback. For the RNA feedback to work as intended, we must ensure that 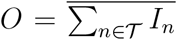, i.e. the logic gate’s output must be inverse to every input targeted by RNA feedback. Hence, because inversion is unambiguous in a Boolean setting, Σ_*n∈*T**_ *I*_*n*_ fully determines the output *O*’s value. As a consequence, any other inputs *n*^*′*^ ∈ *ℐ* with *n*^*′*^ *∉ *T**, i.e. inputs that are not targeted by RNA feedback, necessarily become *don’t care*, i.e. they have no influence on *O*’s value. Removing the inputs that are *don’t care*, the part extended by the RNA feedback must thus be logically inverting, i.e. implement NOR logic on the level of RNA. Consider again the gate architecture from (*11*) in Fig. 3B. There, the logically inverting part is the repression of the transcript that carries the mRNA of the internal TF of the gate by a previous gate’s TF. Because each previous gate’s TF represses a promoter that is placed upstream of an individual copy of the internal TF’s gene, we can express a combinational circuit with this gate architecture as a tree of elementary OR and NOT gates on the molecular level (RNA or Protein) by associating each species with the individual gene copy it originates from. Inclusion of the RNA feedback is then viable. Using a single gene driven by a tandem promoter instead does not allow such a decomposition on the molecular level and our logic requirement 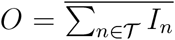 cannot be satisfied. This prohibits the use of our RNA feedback in the NOR gates from (*10*), but allows its use for those from (*11*).

#### 2.4.2 The RNA-Feedback NOR-logic Circuit

Let the topology of a NOR-logic circuit be a directed graph *G ≡* (*V*, *ℰ*), where the vertices (gates, inputs or outputs) *V ≡* {*v*_1_, …, *v*_*K*_} are enumerated by the totally ordered index set *K ≡* {1, …, *K*}. The set *ℰ ⊂ *V* × *V** of directed edges (i.e. wires) in conjunction with the enumeration *K* lets us define incoming *ℐ*_*k*_ *≡* {*m* | (*v*_*m*_ *→ v*_*k*_) *∈ ℰ, m ∈ ℐ}* and outgoing *O*_*k*_ *≡ {m* | (*v*_*k*_ *→ v*_*m*_) *∈ ℰ, m ∈ ℐ}* index sets each for vertex *v*_*k*_. We also define input *U ⊂ ℐ* and output *Y ⊂ ℐ* index sets for the whole circuit. While the topology *G* is obtained in the logic synthesis step of the automated design pipeline, the technology mapping step introduces an injective labelling function *M* : *V →ℒ* that maps each vertex to an element of a library ℒ, i.e. a set of gate parameters from a gate library. Let the mapped gate parameters *M* (*v*_*k*_) = (*µ*_*k*_, *γ*_*k*_, *β*_*k*_, *δ*_*k*_, *n*_*k*_, *κ*_*k*_, *r*_*k*_) *∈ ℒ* be known for each *k* ∈ *K* \ *U*. We then seek to determine the circuit response *α*_*y*_ for *y* ∈ *Y* for given fixed inputs *α*_*u*_ = const. for *u* ∈ *U*. To achieve this, we need the gates’ output promoter activities *α*_*k*_ for each gate *k*. To determine *α*_*k*_, we further need the RNA concentrations *∀l ∈ ℐ*_*k*_ : *m*_*l→k*_ associated with each input of the gate, i.e. each copy of gate *k*’s gene. Note, that in a NOR-logic circuit as in (*10, 11*), we have |*ℐ*_*k*_| ≤ 2 for any *k*. For convenience, we define the total RNA concentration of gene *k* as 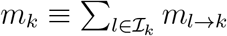. Then, to obtain the circuit response *α*_*y*_ with fixed input *α*_*u*_ = const., we need to solve the set of fixed-point equations

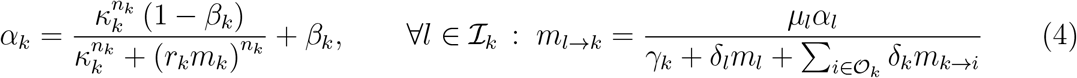

for each *k* ∈ *K* \ *U*. The calculated *m*_*y*_ ∝ [*P*_*y*_] for each *y* ∈ *Y*, proportional to concentrations of reporter proteins [*P*_*y*_], can then be used to score the circuit (*10, 56*).

#### 2.4.3 Cello Circuit End-Stage Example

Translation to Cello’s circuit framework faces several challenges. Cello’s gates feature only parameters *α*_max, *k*_, *α*_min, *k*_, *K*_*k*_ and *n*_*k*_. Matching the parameters from our NOT example in section 2.2 is underdetermined and thus leads to ambiguities. However, we see that *α*_min, *k*_ = *βα*_max, *k*_ and 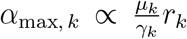 (with any *δ* = 0 we have 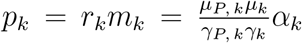.

Although oversimplifying, we can fix the degradation rates *γ*_*P, k*_ and *γ*_*k*_ to values representing their “typical” order of magnitude for *E. coli* (in full knowledge that they may vary greatly in reality). We also set *µ*_*P, k*_ *≡ ω*_*P, k*_ and *µ*_*P, k*_ *≡ α*_max, *k*_*ω*_*k*_ with the proportionality factors *ω*_*P, k*_ and *ω*_*k*_ representing “typical” orders of magnitude for translation and transcription rates. This unifies *r*_*k*_ for all gates and we finally set *κ* = *K*_*k*_*r*_*k*_. With this setup, we took circuit *0×70* from (*10*) and equipped our simulation engine (*57*) with equation system (4) to check how different the circuit behaved with *δ*_*y*_ = 1 for the YFP end-stage. The comparison of the circuit’s closest ON-OFF output pair in the configurations with and without feedback (*δ*_*y*_ = 1 and *δ*_*y*_ = 0) is given in Fig. 3C. More details are found in Methods 4.3.

## 3 Discussion

In this work, we performed a qualitative and quantitative analysis of how an inclusion of a sequestration reaction between the chemical species of an inverting genetic logic gate could impact the dose-response curve of the gate. For this analysis, we chose an asRNA-mediated feedback as the mechanism that implements the sequestration. We pointed out, how the observed effects constitute improvements of the characteristics of the logic gate, that are an increase in steepness of the logic transition region and a reduction in leakage.

The increase in steepness cannot be achieved to an arbitrary degree overall, but numerical analysis has provided evidence that especially in the high-response regime immediately below the inflection point of the original Hill-shaped dose-response curve, strong sequestration can lead to almost vertical sections in the dose-response curve. This qualitative effect has previously been observed (*7*) and is here likely due to the dominance of the hyperbolic feedback term that effectively reduces the concentration of the input RNA and thus the repressing TF. However, as the low-response, high-dose regime is entered, the dose-response curve is dominated by transcriptional repression and the hyperbolic feedforward term, that continually reduce the concentration of output RNA with increasing input promoter activity and thus input RNA concentration.

With an increase in sequestration rate, the dose-response curve can lose injectivity and the logic gate can experience bistability. This effect can lead to a hysteresis in the dose-response curve which can be beneficial in stabilizing logic levels. This hysteresis must be finely calibrated though, as it becoming too strong can cause switching to become almost impossible and it can effectively widen the region in which logic transitions occur. In addition to that, in a genetic logic circuit that consists of a cascade of inverting logic gates, multistability must be very carefully integrated, especially because the logic levels within the circuit are instantiated non-trivially before an inducer is added. Thus, it can be crucial to maintain monostability, which is why we proposed Alg. 1 for parameter tuning.

Leakage reduction is important in many applications, and using RNA sequestration to achieve this is an established concept (*28, 29*). While our model predicts an arbitrarily strong leakage reduction, it certainly misses out on some effects. Those effects may include imperfect blocking of ribosome recruitment by dsRNA formation or spatially heterogeneous sequestration rates due to transport limitations of chemical species. However, especially in scenarios where the choice of possible inducible promoters is limited to such with significant leakage, addition of RNA feedback can be an easy way to achieve strong leakage reduction. The RNA feedback mechanism can be readily adapted to existing logic gates that incorporate inverting logic elements between constituting chemical species, the simplest of those being NOT/NOR gates. Researchers and engineers can integrate this structure into both current and newly developed gates. Tuning gate characteristics to more closely resemble those of an ideal logic gate is important for switches that must avoid intermediate states, such as kill switches. Biosensors can also benefit, as they only need to detect the definite presence or absence of a compound. We gave an outlook on the integration of the RNA-feedback mechanism into genetic circuits with multiple gates. In genetic design automation, integrating RNA-mediated feedback into existing gate libraries is straightforward and can result in enhanced performance in circuits of any size. Circuit simulation and parameter tuning as part of the technology mapping process become more expensive, but time-frames and general cost of wetlab iteration still outweigh drylab computation cost and time by a large margin, justifying the additional expense to be made.

Finally, while the sequestration between chemical species of logic gates has been envisioned to happen in the form of asRNA targeting and dsRNA formation in this work, the specific way this sequestration is implemented is transparent in the theoretical analysis, and other means of implementation are imaginable. Even the use of protein-protein interaction or RNA-protein interaction to implement the sequestration would require only minor changes in the proposed equations and the qualitative statements made w.r.t. the affected species remain.

## 4 Methods

### 4.1 Single NOT Gate Case Study

The parameters for the quantitative study of the single NOT gate presented were chosen to be *µ*_1_ = 1, *µ*_2_ = 10, *γ*_1_ = *γ*_2_ = 0.2, *µ*_*p*_ = 1, *γ*_*p*_ = 0.002, *β* = 10^−4^, *K* = 1 and *n* = 1. For the steady-state analysis using the ODEs, the free parameters *x* and *δ* were chosen to fall in the ranges *x ∈* [10^−4^, 10^2^] and *δ ∈* [0, 1]. Note, that the choice *µ*_1_ = 1 is somewhat arbitrary, because we chose *x* to be a generic input variable. The setup for the stochastic simulation is explained in the following.

#### 4.1.1 Stochastic Simulation of the NOT Gate

The SSA simulation for the NOT gate studied in section 2.2 consisted in total of 8 species and 14 reactions. The two promoters were modelled using a two-state switching leaky promoter model (*58*) comprising of two species *X*_ON_, *X*_OFF_ and *G*_ON_, *G*_OFF_ each with conservation laws *X*_ON_ +*X*_OFF_ = *G*_ON_ +*G*_OFF_ = 1. In our example, we restrict ourselves to a cooperativity index of one (*n* = 1), so that we can model the switching between *G*_ON_ and *G*_OFF_ by simple association-dissociation reactions. The reactions for the switching of the two promoters are then given as

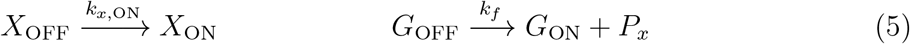

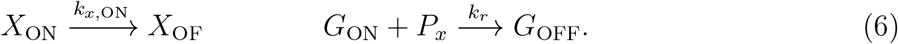

The switching rate constants *k*_*f*_ and *k*_*r*_ were chosen to be consistent with the desired parameters of the black-box (normalized) Hill function of the existing logic gate in (1) and match the time-scale of transcription and translation. Similarly, the switching rates *k*_*x*,ON_, *k*_*x*,OFF_ were chosen to be consistent with the configurable steady-state input promoter activity. The two RNA and two protein species’ *M*_*x*_, *M*_*y*_ and *P*_*x*_, *P*_*y*_ are given modulated birth and death reactions of the form

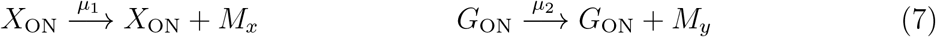

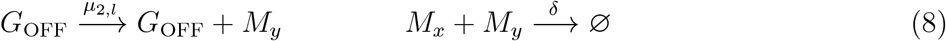

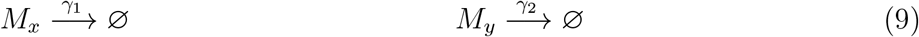

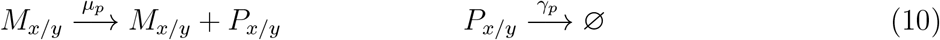

where the slash indicates the same reaction and rate constant is used for both subscripts. The simulations were performed for 10 different input promoter activities *x* = *k*_*x*,ON_(*k*_*x*,ON_ + *k*_*x*,OFF_)^−1^ equally spaced in the log-domain in the interval *x ∈* [10^−2^, 1]. For each input promoter activity, 10 000 trajectories were drawn with sufficient length to reach a steady state. For each obtained steady-state empirical distributions, the median and the 0.1and 0.9-quantiles were calculated and are shown using markers and errorbars in Fig. 2C.

#### 4.1.2 Reaction Rate Equations and Steady State

To derive the reaction rate equations corresponding to the reactions in (5) and (7), we make a quasi-steady-state (QSS) assumption for the association and dissociation events of transcription factors to and from the promoters in the time-scale of transcription and translation (*52*). Thus, we assume

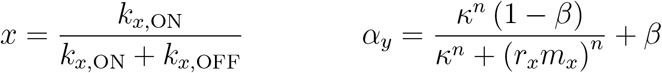

at all times. Because the rate constants *k*_*x*,ON_, *k*_*x*,OFF_ can be freely chosen in this setting, we treat *x* as an independent variable from now on. The reaction rate equations for the remaining species are then given by

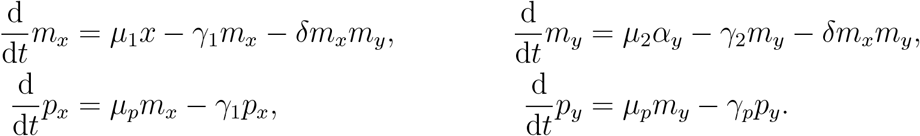

Setting all time-derivatives to zero and rearranging the equations, we obtain the steady-state equations (1).

#### 4.1.3 Uniqueness of the Transition Region

We have shown in section 2.2 that the maximum output RNA concentration *m*_*y*,max_ at *x* = 0 is the same for any choice of sequestration rate *δ*. While the minimum output RNA concentration *m*_*y*,min_ for *x → ∞* is only non-zero in the case of *δ* = 0. However, in the application context of a genetic logic gate, it makes sense to ask for the point *x* = *x*_lv_, where *m*_*y*_(*x*) as a function of *x* crosses a constant line *m*_*y*_(*x*_lv_) = *m*_lv_ that lies in the feasible region of the original NOT gate without feedback, where *δ* = 0, and how often it crosses it, i.e. how many solutions for *x*_lv_ we can find for a given *m*_lv_. Using this rationale, we can not only make statements about the approximate location of the transition region of the extended NOT gate, but we can also determine whether there exist multiple transition regions due to the feedback connection with *δ >* 0 – an undesired property for a logic gate. Consider the steady-state equations (1) from section 2.2. Inserting *m*_lv_ into the equations for *m*_*y*_ and *m*_*x*_, we get

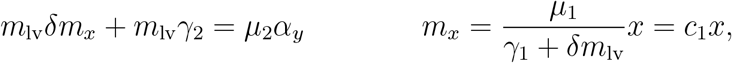

where we introduced the constant *c*_1_ for convenience. We then insert the expression for *α*_*y*_, substitute *m*_*x*_ = *c*_1_*x* and rearrange to get

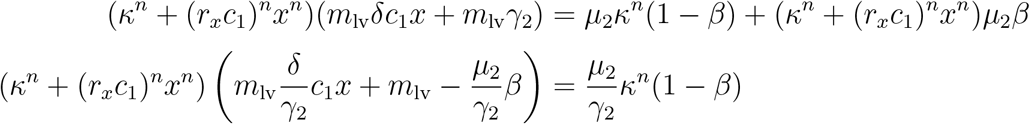

where we divided everything by *γ*_2_ in the last line. At this point, we can recognize 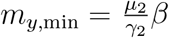 and 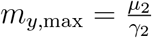 from the original gate *δ* = 0. We insert these and subtract both sides by *κ*^*n*^(*m*_*y*,max_ − *m*_*y*,min_) to get

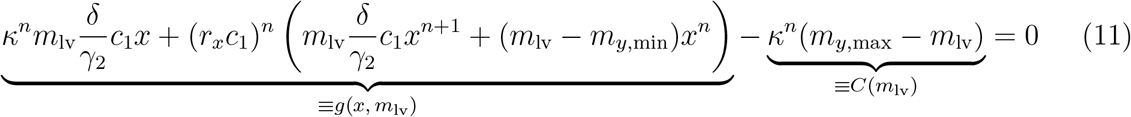

We combine the full LHS of (11) in a single function *f* (*x, m*_lv_) *≡ g*(*x, m*_lv_) − *C*(*m*_lv_). We can now make sense of this equation. If we consider only *m*_lv_ *∈* [*m*_*y*,min_, *m*_*y*,max_], then not only *C*(*m*_lv_) *≥* 0, but also *g*(0, *m*_lv_) − *C*(*m*_lv_) ≤ 0. At the same time, *g* is a (generalized) polynomial in *x* with only non-negative coefficients and thus constitutes a monotonically increasing function for *x ≥* 0, i.e. 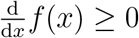. It is in fact strictly monotonically increasing, i.e. 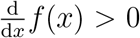, within the interior *x ∈* (0, *∞*) and only violates strict monotonicity at the boundary *x* = 0 for the choice *m*_lv_ = 0. Hence, *f* is bijective in the region *x ≥* 0. Since *f* is also continuous in *x* and (formally) unbounded, *g*(*x, m*_lv_) either has the solution *x* = 0 or changes sign once for *x ∈* (0, *∞*). Thus, *g*(*x*_lv_, *m*_lv_) = 0 has a unique solution *x*_lv_ for any *m*_lv_ *∈* [*m*_*y*,min_, *m*_*y*,max_]. Thus, any concentration *m*_lv_ *∈* [*m*_*y*,min_, *m*_*y*,max_] can only be reached from a unique dose 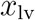.

#### 4.1.4 Reference Location of the Transition Region

Asking for a single point of transition *x*^tr^ akin to the inflection point of a Hill-shaped doseresponse curve might not be answered consistently in general, because the transition region becomes non-symmetrical due to the feedback and is unbounded from below (as can be seen in the example in Fig. 2)C). However, it makes sense to rephrase the question slightly and ask, where the value *m*_*y*_ = *m*_tr_ is reached for *δ >* 0 that constitutes the inflection point of the original NOT gate, where *δ* = 0. The inflection point of the original NOT gate is found at

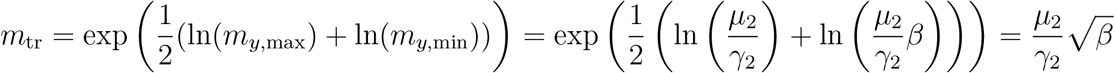

Inserting this in equation (11) gives

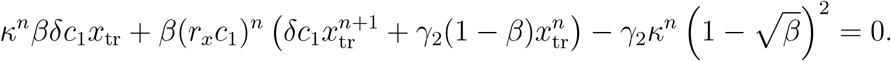

Introduction of the coefficients *d*_1_ *≡ κ*^*n*^*βδc*_1_, *d*_*n*_ *≡ β*(*r*_*x*_*c*_1_)^*n*^(1 − *β*)*γ*_2_, *d*_*n*+1_ = (*r*_*x*_*c*_1_)^*n*^*βδc*_1_ and 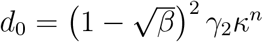 gives eq. (2) from section 2.2.

##### Algorithm 1 Calibration of the Dose-Response Curve

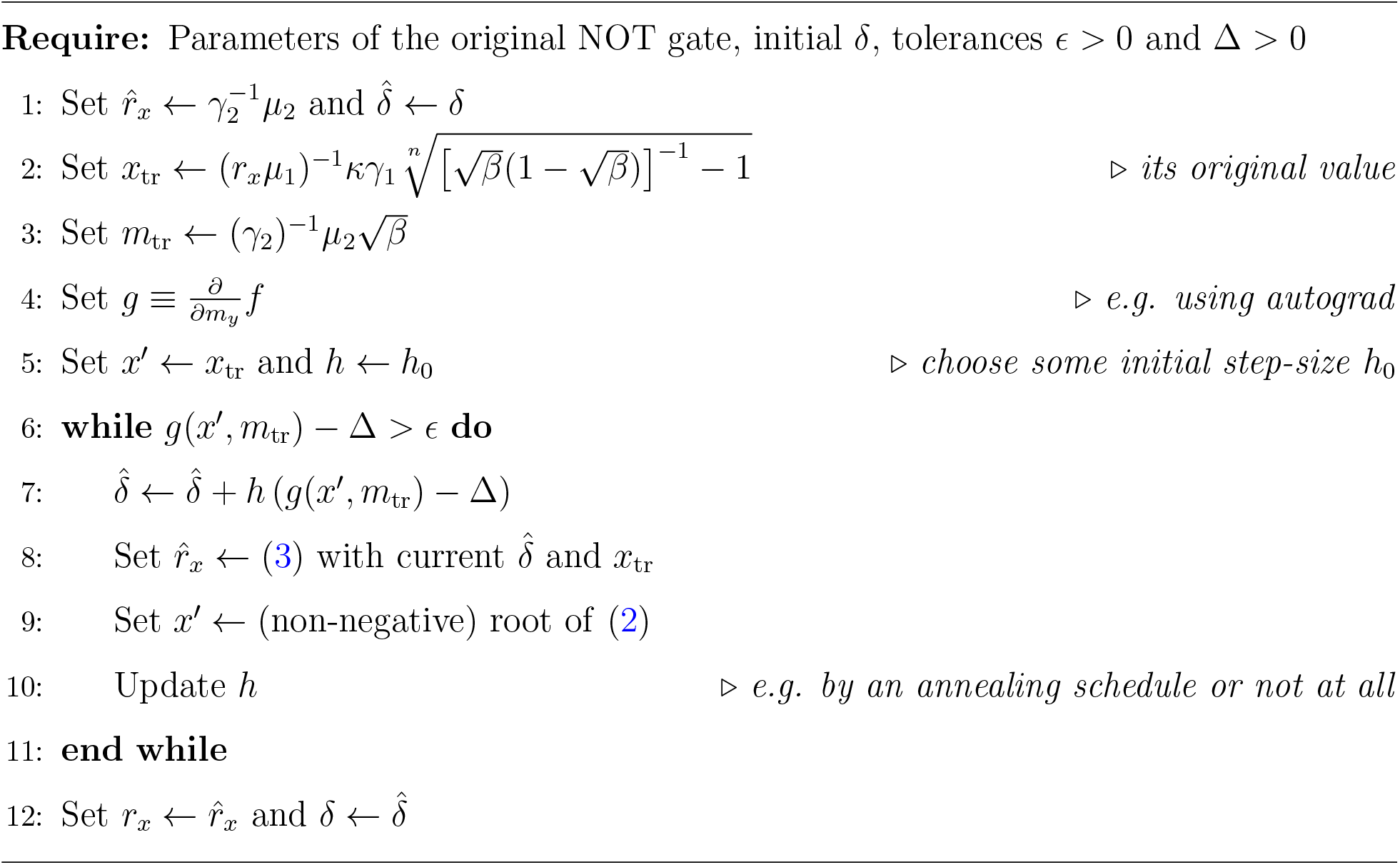

### 4.2 Parameter Tuning Algorithm and Example

We give an exemplary parameter tuning algorithm in Alg. 1. The common design parameters that generated the steady-state curves in the NOT gate example in Fig. 2C are *µ*_1_ = *µ*_2_ = 1, *γ*_1_ = *γ*_2_ = 0.2, *r*_*x*_ = 1, *β* = 10^−2^, *κ* = 0.3 and *n* = 3. In the monostable case, the doseresponse curve coincides with the steady-state curve, and the varying parameters are *r*_*x*_ and *δ*. The vanilla NOT gate’s curve, where there is no RNA feedback, is generated with *r*_*x*_ = 1 and *δ* = 0. The green curve on the LHS of Fig. 2C is generated with *r*_*x*_ = 1 and *δ* = 3 and shows an S-shape typical for a bistable system like the bistable toggle-switch. Application of Alg. 1 then yields the new parameters *r*_*x*_ *≈* 1.22 and *δ* = 0.138 that generate the green curve on the RHS of Fig. 2C, which is monostable and centered around the logarithmic inflection point of the original NOT gate’s dose-response curve.

### 4.3 Circuit Simulation with Feedback using ARCTIC

The circuit simulations in section 2.4.3 are done using the circuit simulator of the CAD software ARCTIC (*57*). ARCTIC uses a fixed-point algorithm to evaluate the circuit response of a logic circuit with feedback connections (that is not combinational) and supports array-valued quantities to be communicated between the logic gates. The latter feature was used to communicate RNA concentrations together with promoter activities between the gates. The simulation library devised for this work is <monospace>libsim iwbda.py</monospace>. The gate library used has been created by fitting the parameters in (4) to the ones of Cello’s gate library (*10*).

## 5 Code Availability

The model is implemented as part of the technology mapping framework ARCTIC, which is available at https://www.rs.tu-darmstadt.de/ARCTIC.

## 6 Author Contributions

N.E. provided mathematical analysis and programming. M.M. provided biological background. H.K. provided the idea. All authors contributed to the writing of this paper.

## 7 Acknowledgments

The authors thank Jérémie Marlhens and Erik Kubaczka for fruitful discussions and proof-reading.

## TOC Graphic

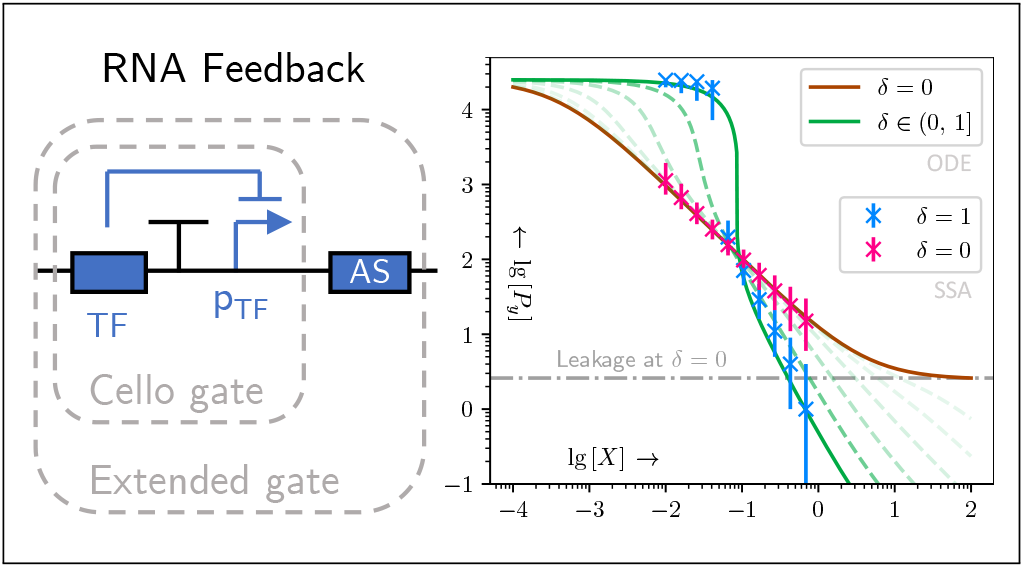

## References

1. Alon, U. An introduction to systems biology: design principles of biological circuits; CRC Press, 2019.

2. Gardner, T. S., Cantor, C. R., and Collins, J. J. (2000) Construction of a genetic toggle switch in Escherichia coli. Nature 403, 339–342.

3. Cardinale, S., and Arkin, A. P. (2012) Contextualizing context for synthetic biology– identifying causes of failure of synthetic biological systems. Biotechnology Journal 7, 856–866.

4. Brewster, R. C., Weinert, F. M., Garcia, H. G., Song, D., Rydenfelt, M., and Phillips, R. (2014) The Transcription Factor Titration Effect Dictates Level of Gene Expression. Cell 156, 1312–1323.

5. Rydenfelt, M., Cox III, R. S., Garcia, H., and Phillips, R. (2014) Statistical mechanical model of coupled transcription from multiple promoters due to transcription factor titration. Physical Review E 89, 012702.

6. Engelmann, N., Schwarz, T., Kubaczka, E., Hochberger, C., and Koeppl, H. (2023) Context-Aware Technology Mapping in Genetic Design Automation. ACS Synthetic Biology 12, 446–459, PMID: 36693176.

7. Jeong, E. M., Song, Y. M., and Kim, J. K. (2022) Combined multiple transcriptional repression mechanisms generate ultrasensitivity and oscillations. Interface Focus 12, 20210084.

8. Brophy, J. A., and Voigt, C. A. (2014) Principles of genetic circuit design. Nature Methods 11, 508–520.

9. Buecherl, L., and Myers, C. J. (2022) Engineering genetic circuits: advancements in genetic design automation tools and standards for synthetic biology. Current Opinion in Microbiology 68, 102155.

10. Nielsen, A. A., Der, B. S., Shin, J., Vaidyanathan, P., Paralanov, V., Strychalski, E. A., Ross, D., Densmore, D., and Voigt, C. A. (2016) Genetic circuit design automation. Science 352, aac7341.

11. Chen, Y., Zhang, S., Young, E. M., Jones, T. S., Densmore, D., and Voigt, C. A. (2020) Genetic circuit design automation for yeast. Nature Microbiology 5, 1349–1360.

12. Xie, M., and Fussenegger, M. (2018) Designing cell function: assembly of synthetic gene circuits for cell biology applications. Nature Reviews Molecular Cell Biology 19, 507–525.

13. Olson, E. J., and Tabor, J. J. (2012) Post-translational tools expand the scope of synthetic biology. Current Opinion in Chemical Biology 16, 300–306.

14. Rosenfeld, N., and Alon, U. (2003) Response delays and the structure of transcription networks. Journal of Molecular Biology 329, 645–654.

15. Bentley, W. E., and Kompala, D. S. (1990) Optimal Induction of Protein Synthesis in Recombinant Bacterial Cultures a. Annals of the New York Academy of Sciences 589, 121–138.

16. Ceroni, F., Algar, R., Stan, G.-B., and Ellis, T. (2015) Quantifying cellular capacity identifies gene expression designs with reduced burden. Nature Methods 12, 415–418.

17. Glick, B. R. (1995) Metabolic load and heterologous gene expression. Biotechnology Advances 13, 247–261.

18. Chappell, J., Takahashi, M. K., Meyer, S., Loughrey, D., Watters, K. E., and Lucks, J. (2013) The centrality of RNA for engineering gene expression. Biotechnology Journal 8, 1379–1395.

19. Schmidt, C. M., and Smolke, C. D. (2019) RNA switches for synthetic biology. Cold Spring Harbor Perspectives in Biology 11, a032532.

20. Dykstra, P. B., Kaplan, M., and Smolke, C. D. (2022) Engineering synthetic RNA devices for cell control. Nature Reviews Genetics 23, 215–228.

21. Lehr, F.-X., Hanst, M., Vogel, M., Kremer, J., Göringer, H. U., Suess, B., and Koeppl, H. (2019) Cell-free prototyping of AND-logic gates based on heterogeneous RNA activators. ACS Synthetic Biology 8, 2163–2173.

22. Qi, L. S., Larson, M. H., Gilbert, L. A., Doudna, J. A., Weissman, J. S., Arkin, A. P., and Lim, W. A. (2013) Repurposing CRISPR as an RNA-guided platform for sequence-specific control of gene expression. Cell 152, 1173–1183.

23. Green, A. A., Silver, P. A., Collins, J. J., and Yin, P. (2014) Toehold switches: de-novodesigned regulators of gene expression. Cell 159, 925–939.

24. Zhao, E. M., Mao, A. S., de Puig, H., Zhang, K., Tippens, N. D., Tan, X., Ran, F. A., Han, I., Nguyen, P. Q., Chory, E. J., and others (2022) RNA-responsive elements for eukaryotic translational control. Nature Biotechnology 40, 539–545.

25. Laurent, G. S., Wahlestedt, C., and Kapranov, P. (2015) The Landscape of long non-coding RNA classification. Trends in Genetics 31, 239–251.

26. Hammond, S. M. (2015) An overview of microRNAs. Advanced Drug Delivery Reviews 87, 3–14.

27. Frei, T., Chang, C.-H., Filo, M., Arampatzis, A., and Khammash, M. (2022) A genetic mammalian proportional–integral feedback control circuit for robust and precise gene regulation. Proceedings of the National Academy of Sciences 119, e2122132119.

28. Meister, G., and Tuschl, T. (2004) Mechanisms of gene silencing by double-stranded RNA. Nature 431, 343–349.

29. Kay, C., Skotte, N., Southwell, A., and Hayden, M. (2014) Personalized gene silencing therapeutics for Huntington disease. Clinical Genetics 86, 29–36.

30. Obbard, D. J., Gordon, K. H., Buck, A. H., and Jiggins, F. M. (2009) The evolution of RNAi as a defence against viruses and transposable elements. Philosophical Transactions of the Royal Society B: Biological Sciences 364, 99–115.

31. Verd, B., Crombach, A., and Jaeger, J. (2014) Classification of transient behaviours in a time-dependent toggle switch model. BMC systems biology 8, 1–19.

32. Bernhart, S. H., Tafer, H., Mückstein, U., Flamm, C., Stadler, P. F., and Hofacker, I. L. (2006) Partition function and base pairing probabilities of RNA heterodimers. Algorithms for Molecular Biology 1, 1–10.

33. Gander, M. W., Vrana, J. D., Voje, W. E., Carothers, J. M., and Klavins, E. (2017) Digital logic circuits in yeast with CRISPR-dCas9 NOR gates. Nature Communications 8, 15459.

34. Tamsir, A., Tabor, J. J., and Voigt, C. A. (2011) Robust multicellular computing using genetically encoded NOR gates and chemical ‘wires’. Nature 469, 212–215.

35. Goñi-Moreno, A., and Amos, M. (2012) A reconfigurable NAND/NOR genetic logic gate. BMC Systems Biology 6, 1–11.

36. Miyamoto, T., Razavi, S., DeRose, R., and Inoue, T. (2013) Synthesizing biomolecule-based Boolean logic gates. ACS Synthetic Biology 2, 72–82.

37. Manzoni, R., Urrios, A., Velazquez-Garcia, S., de Nadal, E., and Posas, F. (2016) Synthetic biology: insights into biological computation. Integrative Biology 8, 518–532.

38. Ferrell, J. E., and Ha, S. H. (2014) Ultrasensitivity part I: Michaelian responses and zero-order ultrasensitivity. Trends in Biochemical Sciences 39, 496–503.

39. Ferrell, J. E., Ha, S. H., and others (2014) Ultrasensitivity part II: multisite phosphorylation, stoichiometric inhibitors, and positive feedback. Trends in Biochemical Sciences 39, 556–569.

40. Ferrell, J. E., and Ha, S. H. (2014) Ultrasensitivity part III: cascades, bistable switches, and oscillators. Trends in Biochemical Sciences 39, 612–618.

41. Jeong, E. M., and Kim, J. K. (2024) A robust ultrasensitive transcriptional switch in noisy cellular environments. npj Systems Biology and Applications 10, 30.

42. Kafri, M., Metzl-Raz, E., Jona, G., and Barkai, N. (2016) The cost of protein production. Cell Reports 14, 22–31.

43. Singh, I. K., Singh, S., Mogilicherla, K., Shukla, J. N., and Palli, S. R. (2017) Comparative analysis of double-stranded RNA degradation and processing in insects. Scientific Reports 7, 17059.

44. Wang, Q., and Carmichael, G. G. (2004) Effects of length and location on the cellular response to double-stranded RNA. Microbiology and Molecular Biology Reviews 68, 432–452.

45. Wery, M., Descrimes, M., Vogt, N., Dallongeville, A.-S., Gautheret, D., and Morillon, A. (2016) Nonsense-mediated decay restricts LncRNA levels in yeast unless blocked by double-stranded RNA structure. Molecular Cell 61, 379–392.

46. Bayer, T. S., and Smolke, C. D. (2005) Programmable ligand-controlled riboregulators of eukaryotic gene expression. Nature Biotechnology 23, 337–343.

47. Briat, C., Gupta, A., and Khammash, M. (2016) Antithetic integral feedback ensures robust perfect adaptation in noisy biomolecular networks. Cell Systems 2, 15–26.

48. Bernstein, J. A., Khodursky, A. B., Lin, P.-H., Lin-Chao, S., and Cohen, S. N. (2002) Global analysis of mRNA decay and abundance in Escherichia coli at single-gene resolution using two-color fluorescent DNA microarrays. Proceedings of the National Academy of Sciences 99, 9697–9702.

49. Selinger, D. W., Saxena, R. M., Cheung, K. J., Church, G. M., and Rosenow, C. (2003) Global RNA half-life analysis in Escherichia coli reveals positional patterns of transcript degradation. Genome Research 13, 216–223.

50. Wagner, E. G. H., Altuvia, S., and Romby, P. (2002) Antisense RNAs in bacteria and their genetic elements. Advances in Genetics 46, 361–398.

51. Nicholson, A. W. (2014) Ribonuclease III mechanisms of double-stranded RNA cleavage. Wiley Interdisciplinary Reviews: RNA 5, 31–48.

52. Bintu, L., Buchler, N. E., Garcia, H. G., Gerland, U., Hwa, T., Kondev, J., and Phillips, R. (2005) Transcriptional regulation by the numbers: models. Current Opinion in Genetics & Development 15, 116–124.

53. Schile, A. J., and Limmer, D. T. (2019) Rate constants in spatially inhomogeneous systems. The Journal of Chemical Physics 150.

54. Mano, M. M., and Ciletti, M. D. Digital Design; Pearson Education, 2002.

55. Steininger, A., Maier, J., and Najvirt, R. The metastable behavior of a schmitt-trigger. 2016 22nd IEEE International Symposium on Asynchronous Circuits and Systems (ASYNC). 2016; pp 57–64.

56. Schladt, T., Engelmann, N., Kubaczka, E., Hochberger, C., and Koeppl, H. (2021) Automated design of robust genetic circuits: Structural variants and parameter uncertainty. ACS Synthetic Biology 10, 3316–3329.

57. Schwarz, K. e. a., Engelmann ARCTIC Design Automation Toolchain. https://gitlab.rs.e-technik.tu-darmstadt.de/arctic/arctic, 2021.

58. Huang, L., Yuan, Z., Liu, P., and Zhou, T. (2015) Effects of promoter leakage on dynamics of gene expression. BMC Systems Biology 9, 1–12.

